# Pectin methylesterase selectively softens the onion epidermal wall yet reduces acid-induced creep

**DOI:** 10.1101/766931

**Authors:** Xuan Wang, Liza Wilson, Daniel J. Cosgrove

## Abstract

De-esterification of homogalacturonan (HG) is thought to stiffen pectin gels and primary cell walls by increasing calcium crosslinking between HG chains. Contrary to this idea, recent studies found that HG de-esterification correlated with reduced stiffness of living tissues, measured by surface indentation. The physical basis of such apparent wall softening is unclear, but possibly involves complex biological responses to HG modification. To assess the direct physical consequences of HG de-esterification on wall mechanics without such complications, we treated isolated onion (*Allium cepa*) epidermal walls with pectin methylesterase (PME) and assessed wall biomechanics with indentation and tensile tests. In nanoindentation assays, PME action softened the wall (reduced the indentation modulus). In tensile force/extension assays, PME increased plasticity, but not elasticity. These softening effects are attributed, at least in part, to increased electrostatic repulsion and swelling of the wall after PME treatment. Despite softening and swelling upon HG de-esterification, PME treatment alone failed to induce cell wall creep. Instead, acid-induced creep, mediated by endogenous α expansin, was reduced. We conclude that HG de-esterification physically softens the onion wall, yet reduces expansin-mediated wall extensibility.

**Highlight:** After enzymatic de-esterification without added calcium, the onion epidermal wall swells and becomes softer, as assessed by nanoindentation and tensile plasticity tests, yet exhibits reduced expansin-mediated creep.

## Introduction

This study attempts to resolve some perplexing and apparently contradictory results concerning the influence of pectin de-esterification on the mechanics and extensibility of growing cell walls. Pectins are acidic polysaccharides that constitute a large proportion of the primary cell wall of many plant species and are particularly dominant in cell walls of some Charophycean algae (Domozych *et al*., 2014) and the tips of pollen tubes (Chebli *et al*., 2012). Homogalacturonan (HG) constitutes the most abundant pectic component of the primary wall (Atmodjo *et al*., 2013). It is synthesized in methyl esterified form in the Golgi system, deposited to the cell wall, and subsequently de-esterified *in muro* by pectin methylesterases (PMEs) (Wolf *et al*., 2009). Methyl esters block the electrostatic charge of galacturonic acid residues that make up the HG backbone and influence the physico-chemical properties of HG, particularly its net charge and charge distribution, its gelling ability, its interactions with cations, notably calcium (Jarvis, 1984; John *et al*., 2019; MacDougall *et al*., 1996; Willats *et al*., 2001) and its interactions with cellulose (Phyo *et al*., 2017).

*In vivo*, HG de-esterification by PME has often been associated with wall stiffening and growth cessation (Goldberg *et al*., 1989; Goldberg *et al*., 1996; Hongo *et al*., 2012; Siedlecka *et al*., 2008). Calcium crosslinking of de-esterified regions of HG is presumed to stiffen the wall, e.g. Bou Daher *et al*. (2018), but quantification of calcium crosslinking is challenging (Hocq *et al*., 2017) and the oft-cited idea that calcium-mediated wall stiffening limits cell wall growth is not well supported, e.g. (Coartney and Morre, 1980; Virk and Cleland, 1990). Moreover, the concept of wall stiffness is not as simple as it once seemed, as Zhang *et al*. (2019) found that indentation stiffness (normal to the plane of the cell wall) can vary independently of tensile stiffness (in the plane of the cell wall) and is not closely linked to wall extensibility, as measured by cell wall creep assays. There is evidence that HG may limit the loosening action of expansins as cells cease growth (Wei *et al*., 2010; Zhao *et al*., 2008), but whether this results from wall stiffening or from interference with expansin action by another means is uncertain. In the special case of pollen tubes, regions of low HG esterification on the flanks of the tip coincide with cell wall regions of high stiffness and reduced surface enlargement, potentially connected with calcium crosslinking of HG (Chebli *et al*., 2012; Zerzour *et al*., 2009).

Contrary to these ideas, other studies reported an opposite pattern in which regions of low HG esterification at the surface of *Arabidopsis thaliana* shoot apical meristems coincided with regions of cell expansion and low mechanical stiffness, as measured by surface micro-indentation with a 5-μm bead (Braybrook and Peaucelle, 2013a; Peaucelle *et al*., 2011; Peaucelle *et al*., 2008). A later study assessing the effect of PME activity on Arabidopsis hypocotyls reported a similar trend (Peaucelle *et al*., 2015). The basis for reduced stiffness in regions of reduced esterification, where high stiffness would generally be expected, is uncertain. Various possibilities have been proposed, but not yet tested: (a) with limited Ca^2+^ supply, de-esterified HGs may not become ionically crosslinked, instead resulting in a more fluid HG; (b) de-esterified HGs may be degraded by endogenous endo-polygalacturonase (PG) or pectate lyase (PL), thereby becoming more fluid; and (c) PME activity may acidify the wall, potentially activating expansins or weakening direct physical interactions between pectin and cellulose, e.g. Phyo *et al*. (2019). In addition to these biochemical possibilities, HG modifications *in vivo* may trigger complex biological responses involving wall integrity sensors, brassinosteroids, auxin and other signaling pathways, with undefined consequences for cell wall properties (Braybrook and Peaucelle, 2013b; Wolf *et al*., 2012). Reviews of this topic point out the many complications involved in relating growth and wall mechanics to HG methylesterification (Levesque-Tremblay *et al*., 2015; Peaucelle *et al*., 2012). These considerations highlight the difficulty and uncertainty in ascribing a direct causal relationship between pectin esterification and wall mechanics in growing organs.

With these points in mind, we designed an *in-vitro* experimental approach to identify the direct consequences of PME action on wall mechanics and extensibility, without the inherent complications and secondary responses likely in living cells. Although *in-vitro* experiments have demonstrated Ca^2+^-mediated stiffening of pectic gels by PME (Willats *et al*., 2001), there appears to be scant information about the direct (physical) consequences of PME action on cell wall biomechanics.

For this study we used the cell-free outer periclinal wall from the onion scale epidermis, because it lends itself to nano-scale indentation and macro-scale tensile tests (Zhang *et al*., 2019; Zhang *et al*., 2017). The outer epidermal wall represents a major structural restraint to growth of many organs (Cosgrove, 2018b; Galletti *et al*., 2016; Kutschera, 1992; Kutschera and Niklas, 2007) and epidermal wall stresses may participate in feedback loops that modulate microtubule patterns and morphogenesis, e.g. Verger *et al*. (2018). Because cell walls are multilayered, anisotropic structures, we probed PME effects on wall biomechanics with both tensile and indentation assays, recognizing that changes in these distinctive mechanical properties may not be closely coupled (Zhang *et al*., 2019).

As described elsewhere (Cosgrove, 2018a), we make a distinction between wall softening and wall loosening: ‘softening’ makes the wall more deformable to mechanical force (a purely mechanical concept) whereas ‘loosening’ induces wall stress relaxation, leading to irreversible wall enlargement, an essential aspect of cell growth and morphogenesis. Zhang *et al*. (2019) found that enzymatically induced wall softening was not sufficient to induce wall loosening.

Loosening is conveniently measured by chemorheological creep of a cell wall (slow, irreversible extension that depends on wall modifying agents such as expansin) whereas softening is measured with rapid force/extension assays that assess wall stiffness. In their simplest forms, indentation assays measure out-of-plane wall stiffness while tensile assays measure in-plane stiffness. As shown below, PME treatment indeed softens the wall in some (but not all) respects, yet does not result in wall loosening and in fact reduces the loosening action of endogenous expansins.

## Materials and methods

Distilled/de-ionized water (18 megohm-cm) was used throughout. Chemicals and reagents were analytical grade. Suppliers for enzymes and antibodies are listed below.

### Cell wall preparation

White onion bulbs (*Allium cepa*), ∼15 cm in diameter, were purchased from local grocery stores. The 5th scale, with the 1st being the outermost fleshy scale, was used to make epidermal peels. Abaxial epidermal cells were torn open by peeling 3-mm or 5-mm wide strips midway along the apical-to-basal gradient of the scale, as described previously (Zhang *et al*., 2014; Zhang *et al*., 2019). The epidermal peels were washed with 20 mM HEPES (4-(2-hydroxyethyl)-1-piperazineethanesulfonic acid), pH 7.5, with 0.01% (v/v) Tween 20 for 15 min to eliminate residual cytoplasmic debris, then dipped in boiling methanol for 30 s to deactivate endogenous wall enzymes while retaining α-expansin activity (Cosgrove and Durachko, 1994).

### Pectin methylesterase

Recombinant PME (Uniprot accession #Q829N4) from *Streptomyces avermitilis* (Cat. #PRO-E0233, 27.5 units/mg; PROZOMIX, Haltwhistle, UK) was desalted with 3 kDa centrifugal filters (Merck Millipore, Tullagreen, IRL) and diluted to 50μg/mL in 20mM HEPES, pH 7.5, for all experiments except where noted. The supplier indicates this PME is a processive enzyme with an activity maximum at pH 8.5, reduced to 70% of maximal activity at pH 7.5.

### Antibody labeling

A strip of onion epidermal wall (10 mm × 10 mm) was placed onto a glass slide with the inner surface facing upward and affixed by sealing the edges with nail polish. The exposed epidermal wall inner surface was submerged in 20 mM HEPES pH 7.5 ± 50μg/mL PME for 2 h at room temperature. The samples were then washed with 1× PBS (phosphate-buffered saline) three times. To block the wall, 1× TBS (tris-buffered saline) based blocking agent containing 10% (w/v) horse serum, 2 mM sodium azide, and 0.01% (v/v) Tween 20 was dropped onto the exposed wall surface for 1 h. JIM7 or LM19 antibodies (PlantProbes, Leeds, UK), diluted 10×, were bound to the wall surface for 1 h, followed by addition of 100× diluted secondary antibody: FITC-linked anti-rat IgG (Thermo Fisher Scientific, Rockford, IL) for an additional 1 h. Samples were washed extensively with 1× PBS three times at the end of each antibody labeling step. Labeled wall samples were imaged with an Olympus BX63 microscope using the FITC channel λex = 490 nm, λem = 525 nm).

### Quantification of methanol release by saponification and PME

Methanol quantification was based on the alcohol oxidase method (Klavons and Bennett, 1986). Onion wall strips (3 mm × 10 mm) were peeled, washed with 20 mM HEPES pH 7.5 with 0.01% (v/v) Tween 20 for 15 min, and boiled in water for 10 s to inactivate endogenous PME and other wall enzymes. Three wall strips were incubated at room temperature in 500 μL 20 mM HEPES pH, 7.5, containing 50 μg/mL PME for 0.5, 1, 1.5, 3, 6, and 16 hours. A negative control was prepared by incubating heat-inactivated wall strips in buffer for 16 h. For quantifying the total saponifiable methyl esters in the wall, three wall strips were placed in 500 µL 1 M NaOH for 1 h. The supernatant was collected from each sample and filtered through a 0.4 μm centrifugation filter. For the NaOH saponified samples, 10 M HCl was used to adjust the pH to 7.5. Alcohol oxidase (# A2404, Sigma Aldrich, Saint Louis, MO) was added in the amount of 0.03 U to the filtered supernatant and the volume was adjusted to 1 mL with 20 mM HEPES pH 7.5. The mixture was incubated at 26°C for 15 min. A total of 500 µL assay solution (20 mM acetyl acetone, 50 mM acetic acid, and 2 M ammonium acetate) was then added to the reaction followed by incubation at 60°C for 15 min. The reaction was cooled to room temperature before the absorbance was assessed at 412 nm. A standard curve was generated using 1, 2, 4, 8, and 16 µg methanol.

### AFM imaging and nanoindentation

Onion epidermal walls were fixed onto a glass slide by nail polish and the exposed wall surface was immersed in 20 mM HEPES pH 7.5. AFM topography images were captured with a Dimension Icon AFM (Bruker, CA, USA) with a ScanAsyst-Fluid+ probe (nominal spring constant: 0.7 N/m). The tip of the silicon nitride probe has a triangular pyramid shape, 2.5 – 8.0 µm in height with a 15±2.5° front angle, 25±2.5° back angle and 2 nm tip radius. The AFM was operated in PeakForce Tapping mode with ScanAsyst and quantitative nanomechanical mapping.

For nanoindentation, wall samples were affixed by double-sided tape on a glass slide and the edges were sealed with nail polish (Zhang *et al*., 2014). Walls were incubated in 20 mM HEPES pH 7.5 buffer and a series of nanoindentations were carried out with the AFM described above. The deflection sensitivity of the AFM tip was obtained by tapping onto the glass slide (Butt *et al*., 2005). The spring constant of the tip was then calibrated using a thermal tune method (Hutter and Bechhoefer, 1993). A target force of 4 nN was used for the nanoindentation experiments with the ramp size set at 500 nm and ramp speed of 3 μm/s. Five spots (100 nm apart on a straight line parallel to the cell long axis) per cell were chosen for indentation measurements using the Autoramp function in the Nanoscope software (Bruker). For each wall sample, 10 cells at random locations were used. Wall samples were then incubated at room temperature in HEPES buffer ± 50 µg/mL PME on the AFM stage for 3 h and indentations were conducted again for the same X-Y coordinates. The indentation modulus was calculated using the 10% − 90% region of the force curve and the Sneddon model (Sneddon, 1965) in the Nanoscope Analysis program (Bruker).

### Tensile test (force/extension)

Onion wall strips (3 mm × 10 mm) were prepared the same way as in the methanol release experiment. Three strips were placed in a 2 mL centrifuge tube with addition of 500 µL 20 mM HEPES pH 7.5 buffer. For PME treatment, walls were treated with 50 µg/mL PME in HEPES buffer on a shaking block at 26°C, 500 rpm. Tensile tests were conducted on a custom extensometer (Durachko *et al*., 2017). Preliminary assays showed that walls given a 16-h PME pretreatment showed more consistent results than shorter incubations, therefore the 16-h treatment was used. To assess cation effects on wall tensile compliances, PME-treated and control walls were incubated in 100 mM CaCl_2_ or MgCl_2_ for two h before measurement. This rather high concentration was selected to be consistent with previous experiments (Xi *et al*., 2015).

Wall strips were clamped at both ends with two clamps that were 3 mm apart, then extended at 3 mm/min until a load of 10 g (0.1 N) was reached, then returned to the original position. Wall samples were extended a second time to the same load to assess the elastic properties of wall samples (the second load cycle is reversible; see Supplemental Fig. 2 in Zhang *et al*. (2017)). A least-squares fit to the last 10% of each recorded force/extension curve was used to calculate the total and elastic compliances, with the plastic compliance calculated as the difference (Durachko *et al*., 2017). Compliance is the reciprocal of material stiffness (which is proportional to modulus).

**Fig. 1.**
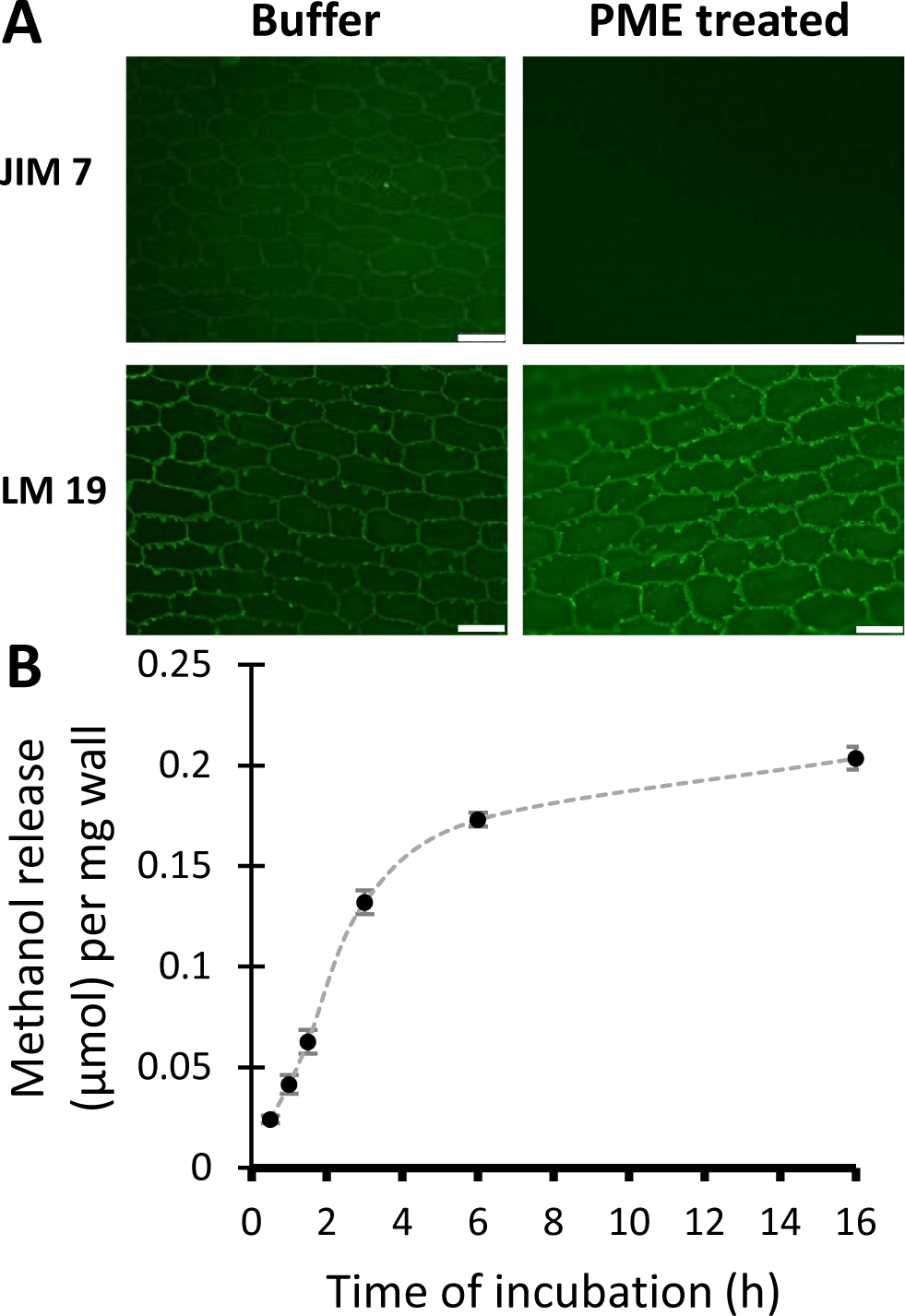
De-esterification of HG in onion epidermal walls by PME. **(A)** Immunolabelling of the inner surface of onion epidermal cell walls after 2 h in HEPES buffer ± PME. JIM7 and LM19 antibodies detect HG of high and low esterifcation, respectively. Paired images ± PME were captured with identical light intensities and camera exposure settings. Scale bar = 100 µm. sRepresentative results from two independent replicates. (**B**) Methanol release from onion epidermal wall as a function of PME treatment time. Mean ± SEM of two independent replicates).

**Fig. 2.**
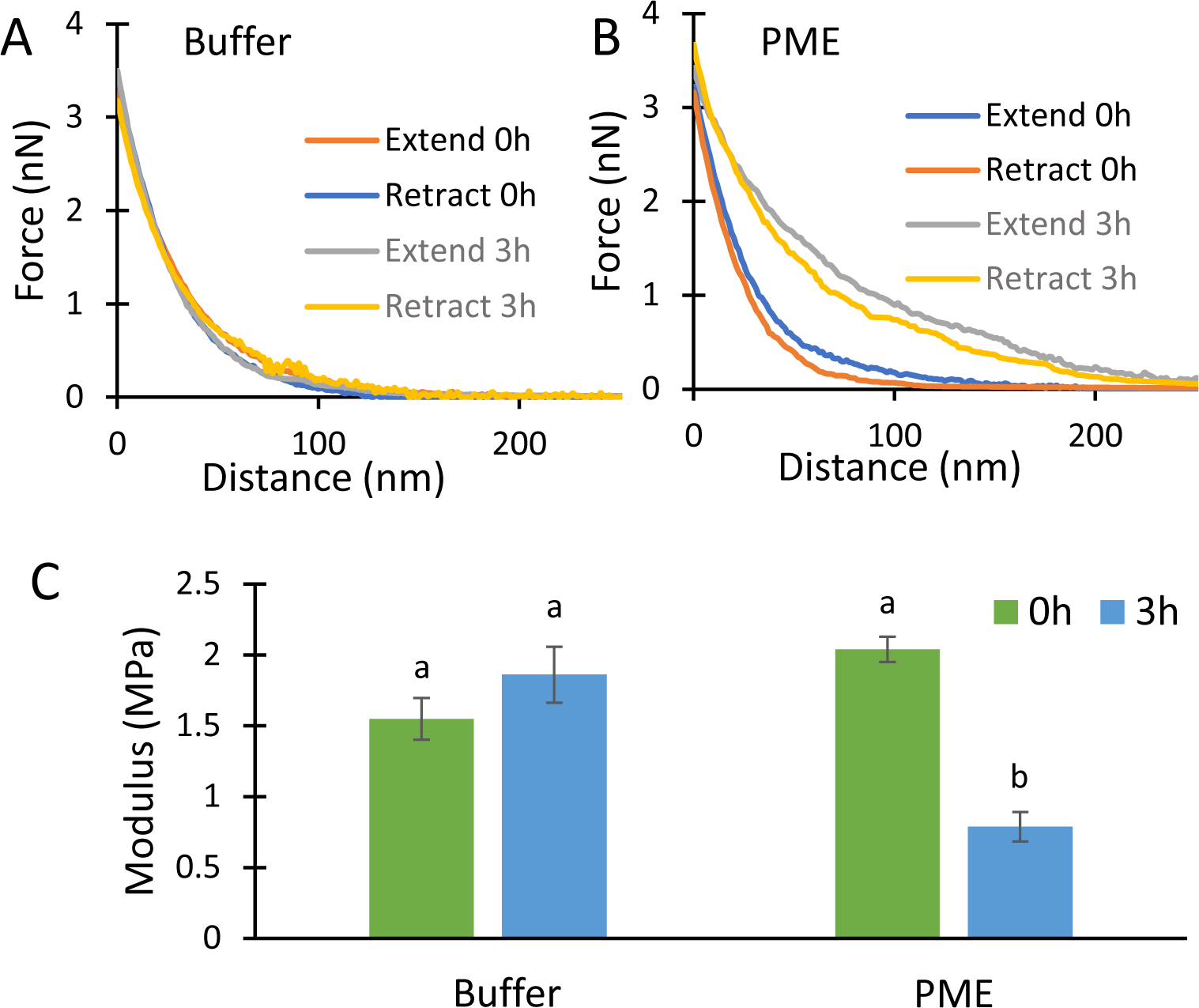
PME effect on AFM nanoindentation. Force-indentation measurements performed on onion epidermal walls submerged in 20 mM pH 7.5 HEPES buffer alone (**A**) and in buffer containing 50 µg/mL PME (**B**) at time 0 h and 3 h. The representative extend and retract curves have modulus values close to the average value of each treatment. (**C**) Modulus calculated by the Sneddon model using retract curves. Values are mean ± SEM (23 ≤ n ≤ 24). Letters indicate statistical significance based on one-way ANOVA with *post hoc* Tukey test (*p* < 0.01).

### Creep and stress relaxation

Creep experiments were carried out at 26°C by clamping wall strips (3 × 10 mm; 5 mm between clamps) in a cuvette filled with buffer, applying a constant tensile force with a 10 g weight (0.1 N) and monitoring specimen length (Durachko *et al*., 2017). To assess PME’s effects on endogenous acid-induced creep, methanol-boiled wall samples were pre-incubated in HEPES pH 7.5 ± 50 μg/mL PME for 16 h at 26°C. Methanol boiling inactivates endogenous enzymes but not expansins. To measure acid-induced extension, walls were clamped in neutral buffer [20 mM MES (2-(N-morpholino) ethanesulfonic acid), pH 6.8, with 2 mM dithiothreitol (DTT)]. When the extension rate became constant, the buffer was replaced with acidic buffer (20 mM NaOAC, pH 4.5, containing 2 mM DTT) to induce expansin-mediated extension. DTT is added to stabilize expansin activity.

To test the ability of PME alone to induce cell wall extension, methanol-boiled walls were clamped in the extensometer in 20 mM HEPES pH 7.5 buffer, as described above. After stabilization of the creep rate, the buffer was exchanged for the same buffer containing 100 µg/mL PME.

To test the effect of buffer pH on wall stress relaxation, 5 mm × 10 mm wall strips were briefly pre-incubated in 20 mM MES pH 6.8 or NaOAC pH 4.5 buffers, with 2 mM DTT, then clamped a custom extensometer with 5 mm between clamps and extended to 120% of their original length, corresponding to a 30 g (0.3 N) load. The decay in holding force was then monitored for 5 min. A small droplet of buffer was positioned at the base of the wall sample to maintain hydration of the sample during the measurement, which was carried out at 26°C. The force decay curves were smoothed using Origin 9.1 (OriginLab, Northampton, MA) and the relaxation rate was calculated by taking the derivative of stress decay with respect to log_10_ time. To test the effect of PME on stress relaxation, wall samples were pre-incubated in 20 mM HEPES buffer pH 7.5±μg/mL PME for 3 h at 26°C, then clamped and stretched as detailed above.

### Onion scale cross-section

A tissue piece (3 mm × 3 mm × 10 mm) was cut from the 5th scale with the longer edge parallel to the long axis of the cells. The onion piece was then placed under a dissecting microscope with the abaxial side facing upward. A razor blade was aligned parallel with the shorter edge in order to make 0.4-mm-thick cross sections by hand. Sections were boiled in methanol for 30 s and stained with 0.1 % toluidine blue (in 20 mM HEPES pH 7.5) for 1 min followed by water washes until the rinses appeared colorless. One section was placed on a glass slide and an O-ring shaped piece of double-sided tape was used for spacing between the glass slide and cover slip. Images of wall cross sections were acquired after 0 h and 3 h incubation in HEPES buffer ± using a 20× objective by an Olympus X63 microscope with an XM10 digital camera.

### Zeta potential measurements

Onion epidermal walls were pulverized at 30 Hz shaking for 10 min (Retsch Cryomill) to produce wall fragments (∼10 µm in diameter) suitable for zeta-potential measurements. Cell walls were prepared as alcohol-insoluble residues following the protocol of Pettolino *et al*. (2012). In brief, wall fragments were incubated in 70% ethanol for 30 min followed by two washes in chloroform:methanol (1:1) for 5 min and one wash in absolute acetone (5 min). Wall fragments were then extensively washed with water; fragments that remained in the suspension after 15 min of settling were collected by filtration and freeze dried. One mg of wall was resuspended in 1 mL buffer and incubated for 16 h at 26°C with 50 µg/mL PME or with 50 µg/mL bovine serum albumin (BSA), a basic protein without wall-specific activity (=negative control). A Zetasizer (Malvern, UK) was used to assess cell wall zeta-potential.

### Image analysis for surface roughness measurement

AFM height images of onion wall surface were flattened (sixth order fit) using the Nanoscope Analysis program (Bruker) to correct for image tilting and large-scale uneveness. Surface roughness was estimated using the built-in measurement of Nanoscope Analysis program with the zero crossing enabled under peak characterization (Longuet-Higgins, 1957). Since the PME treatment reduced the visibility of cellulose microfibrils as imaged in peak force error maps, we anticipated that the peak density (density of detected edges of cellulose microfibrils) would decrease after PME treatment.

## Results

For our initial experiments we prepared cell-free strips of onion epidermal walls (Zhang *et al*., 2019) for comparative analysis by nanoindentation, tensile force/extension testing, and creep testing. Suitably prepared walls were pretreated with buffer ± PME to assess PME’s effects on wall properties as detected by these different biomechanical assays. The first step was to establish appropriate PME treatment.

### De-esterification by PME

Wall strips were incubated for 2 h in 20 mM HEPES buffer, pH 7.5, ± 50 µg/mL PME, which hydrolyzes methyl esters of HG. PME effectiveness was assessed qualitatively by immunofluorescence microscopy with monoclonal antibodies JIM7 and LM19 to detect HG of high and low degree of methylesterification, respectively (Verhertbruggen *et al*., 2009). For buffer-treated walls, labeling by JIM7 and LM19 was most intense for the torn anticlinal walls and the cell borders, with signs of weaker, diffuse labeling of the cell wall proper (Fig. 1A). The labeling pattern indicates the epidermal wall contains a mixture of HG of high and low esterification. After PME treatment, the JIM7 signals were reduced whereas LM19 signals increased, confirming HG de-esterification by PME. Because PME action may alter the accessibility of immunoprobes to their epitopes, we regard these results merely as qualitative evidence that PME treatment indeed modified the wall.

To quantitatively assess the rate and extend of de-esterification, PME-catalyzed release of methanol was measured as a function of time (Fig. 1B). PME acted rapidly and approximately linearly for the first three hours, after which the rate slowed down and approached a plateau after the sixth hour. Approximately 42% of the total saponifiable methyl ester was released during PME treatment for 16 h. Based on this time course, the effect of a 3-h PME treatment, releasing ∼27% of the total saponifiable methyl ester in the wall, was tested in wall biomechanical assays. We estimate that this treatment resulted in larger changes in wall esterification than those that occur as young growing cell walls become mature, e.g. (Goldberg *et al*., 1989; Phyo *et al*., 2017); hence the treatment may represent a relatively large change relative to most biological contexts.

### PME softens the wall in nanoindentation assays

Wall strips were placed cuticle side down on the AFM stage, submerged in buffer, and the upper surface was indented with a sharp pyramidal AFM probe (nominal tip radius 2 nm). This procedure indented the most recently-deposited cell wall surface. The tip pressed ∼120-150 nm into the wall, corresponding to the depth of 3-4 lamellae (Zhang *et al*., 2016), before reaching the target force of 4 nN (Fig. 2A). There was little hysteresis between the indent and retract curves, indicating predominantly elastic indentation with little viscoelastic dissipation.

After 3-h incubation in buffer ± PME, a second set of indentations were made (Fig. 2A, B). The force/distance curves showed the PME-treated walls to be substantially softer than buffer controls. Wall stiffness was quantified as an indentation modulus calculated by the Sneddon model, as appropriate for a sharp pyramid-shaped probe. Because of uncertainties in the shape of wall deformation during indentation and the nonlinear mechanics of cell walls, we view the calculated modulus as an ad-hoc estimate of wall indentation stiffness rather than an accurate estimate of the wall modulus. Buffer-treated walls became slightly stiffer after 3 h, but the difference was not statistically significant (Fig. 2C). In contrast, the resistance to indentation after PME treatment was reduced by 65% in this experiment, as judged by the modulus value.

These results revealed a wall-softening action by PME as detected by nanoindentation. Note that external calcium was not included in the buffer, so this represents the direct effect of PME in a calcium-limited situation. This result supports speculation that PME action might increase HG fluidity when calcium was limiting (Peaucelle *et al*., 2008). Calcium addition stiffened both untreated and PME-treated walls (∼2X and ∼5X, respectively; Supplemental Tables S1-2), consistent with previous work showing that addition or removal of calcium increased or decreased, respectively, the indentation stiffness of untreated onion epidermal walls (Xi *et al*., 2015; Zhang *et al*., 2014; Zhang *et al*., 2019).

### PME potentiates macro-scale plastic deformation in tensile testing

A custom extensometer was used to assess PME’s effect on wall tensile stiffness. Following a conventional protocol (Durachko *et al*., 2017), onion wall strips were stretched and relaxed twice in succession. The second extension is reversible and the slope of the last 10% of the curve is used to calculate an elastic compliance (=slope of the line). Compliance is inversely related to stiffness and modulus. Because the shape of the force/extension curve is nonlinear, compliance values vary with strain. The first extension combines elastic and plastic extensions and is used to calculate a total compliance. A plastic compliance is obtained as the difference between total and elastic compliances. With 0.1 N tensile force, buffer-treated wall strips extended ∼9.5% whereas after PME treatment the extension was ∼12% (Fig. 3A). The additional extension of the PME-treated walls resulted from an increased plastic compliance, whereas the elastic compliance was not significantly affected (Fig. 3B). These results show that PME softens the onion wall in a selective manner, increasing plastic but not elastic deformation.

**Fig. 3.**
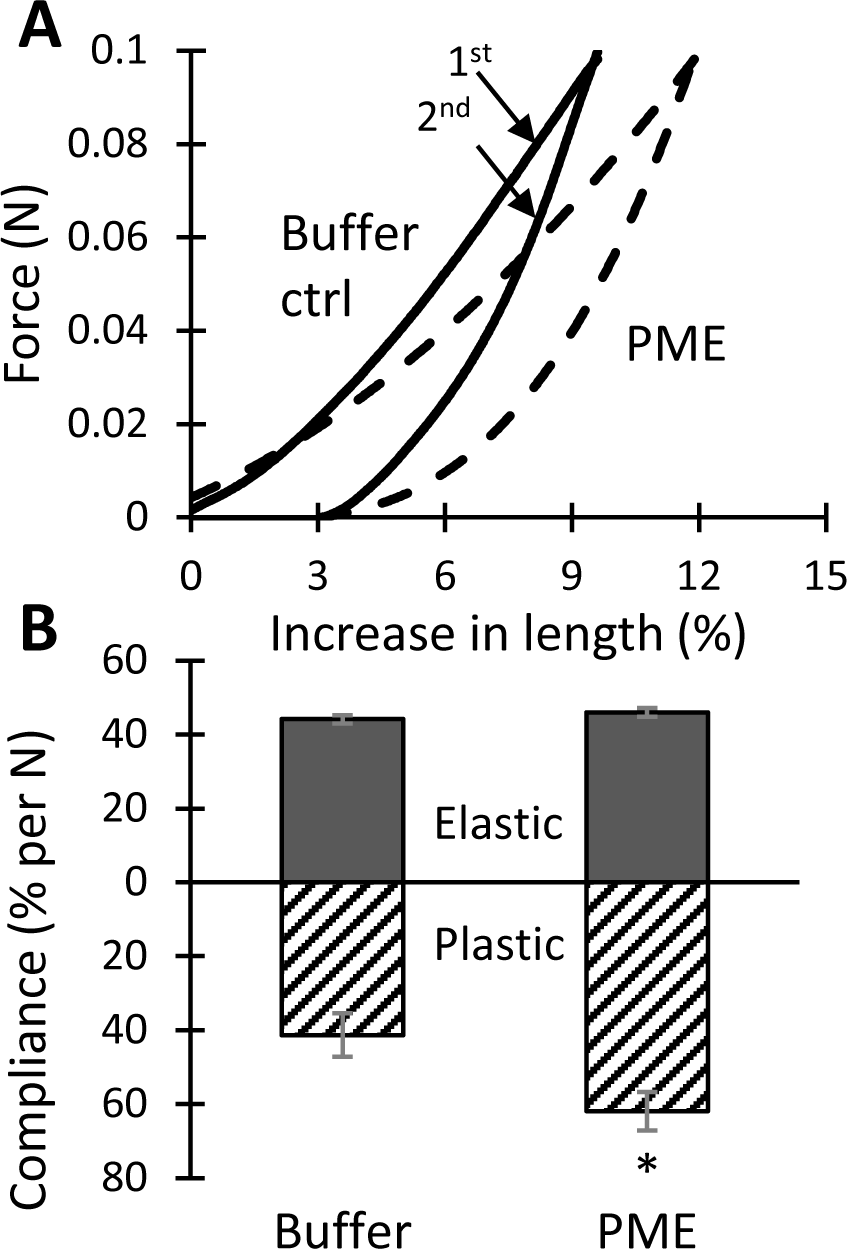
PME effect on tensile mechanics of onion epidermal walls. (A) Force/extension curves of control (solid) and PME-treated (dashed) walls. For each data set, the first stretch is the upper curve which contains both plastic and elastic components while the second stretch is the bottom curve containing only the elastic component. These are representative curves with compliance values close to the average. (B) Statistical summary of elastic and plastic compliances (mean ± SEM; n = 15). Student’s t-test (paired, two-tail) was used to assess statistical significance (* *p* < 0.05). Experiment was repeated four times with similar results.

In these experiments, exogenous calcium was not added to the walls, so enhanced calcium crosslinking of HG after PME treatment was unlikely. We found that addition of 100 mM CaCl_2_ had remarkably little effect on the elastic or plastic compliances of buffer-treated walls (Fig. 4), whereas after PME pretreatment the plastic compliance was reduced by 32% (Fig. 4). This rather high concentration of calcium was used to match that of a previous study on onion indentation (Xi *et al*., 2015) and to be certain that calcium binding sites would be completely saturated.

**Fig. 4.**
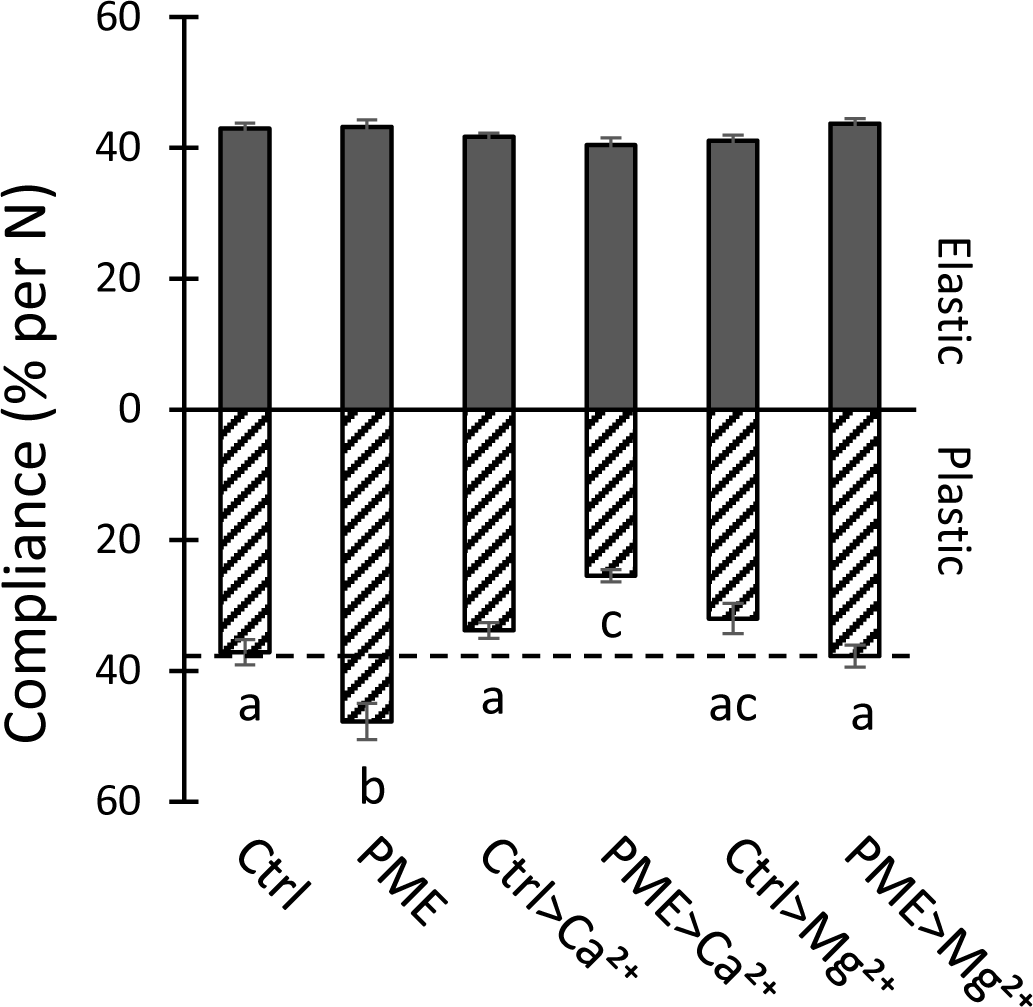
Effect of 100 mM CaCl_2_ and MgCl_2_ on elastic and plastic compliances of walls pretreated with buffer ± PME. Values are means ± SEM (9 ≤ n ≤ 17). Letters indicate statistical difference in one-way ANOVA with *post hoc* Tukey test (*p* < 0.05). Replicated three times for MgCl_2_ and twice for CaCl_2_.

Thus, HG de-esterification reduced wall plasticity in the presence of abundant Ca^2+^, increased plasticity without added Ca^2+^, and had no effect on the elastic compliance in either case. The lack of effect on tensile elasticity is particularly notable.

We also tested the effect of 100 mM MgCl_2_ because Mg^2+^ reportedly does not form strong ionic crosslinks with HG, yet can shield negative charges on the HG backbone (Thibault and Rinaudo, 1986). Mg^2+^ slightly reduced the plastic compliance of buffer-treated walls (not statistically significant) and completely negated PME-mediated softening (Fig. 4). In light of these results we also tested Mg^2+^ in the indentation assay (Supplemental Tables S3-4) where we observed a similar effect: Mg^2+^ suppressed the PME-induced reduction in the indentation modulus, yet had no significant effect on indentation of untreated walls. These results suggest that PME-mediated softening depends, at least in part, on the increased negative electrostatic charge of de-esterified HG and that suppression of PME-enhanced electrostatic charge with high cation concentrations suppresses this softening.

Consistent with this interpretation, zeta potential measurements confirmed that PME-treated walls have a more negative zeta potential than control walls (−23.3 ± 0.41 mV for PME treated versus −17.0 ± 0.35 mV for BSA control, Fig. 5). Thus, PME treatment had two opposing effects: by itself it increased the plastic compliance and reduced the indentation modulus, and it sensitized the wall to exogenous Ca^2+^, exaggerating the stiffening effects of that cation.

**Fig. 5.**
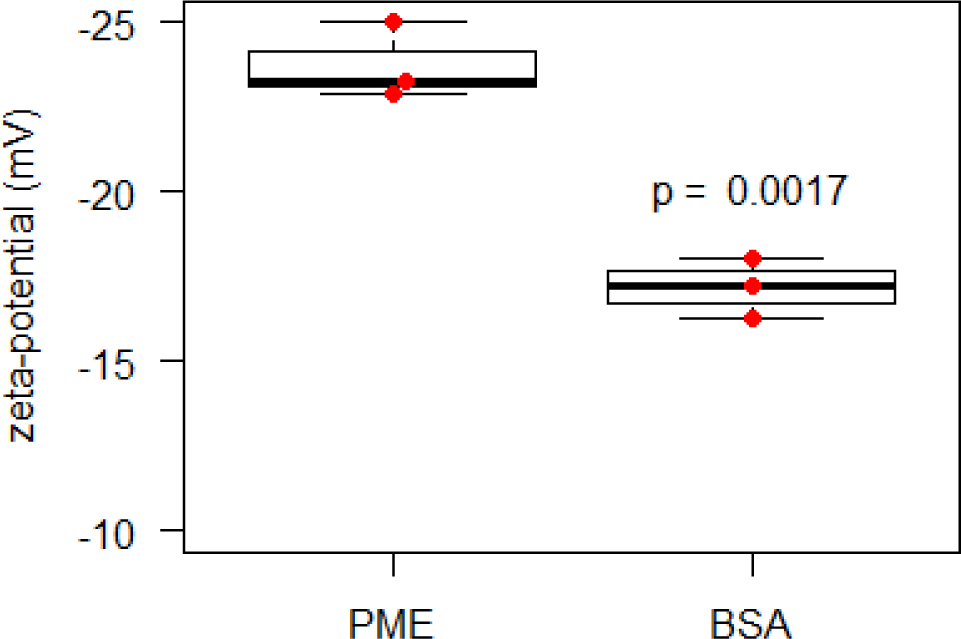
Zeta potential of onion wall fragments treated with PME or BSA (box and whiskers plot). Wall fragments were suspended in 20 mM HEPES pH 7.0. Three biological samples with statistical analysis by Student’s t-test.

### Cell wall swelling upon de-esterification

Electrostatic repulsion between pectin carboxylates generated by PME action could lead to increased hydration and swelling of the cell wall (Ryden *et al*., 2000). In light of this idea, we tested whether PME treatment caused onion wall swelling, assessed microscopically as epidermal wall thickness (Fig. 6A). In these experiments the average thickness of onion epidermal wall increased from 6.2 ± 0.3 µm in buffer to 7.3 ± 0.2 µm after 3-h incubation with PME (Fig. 6B), whereas incubation in buffer alone did not alter wall thickness significantly. Thus, we conclude that HG de-esterification indeed enhanced wall hydration.

**Fig. 6.**
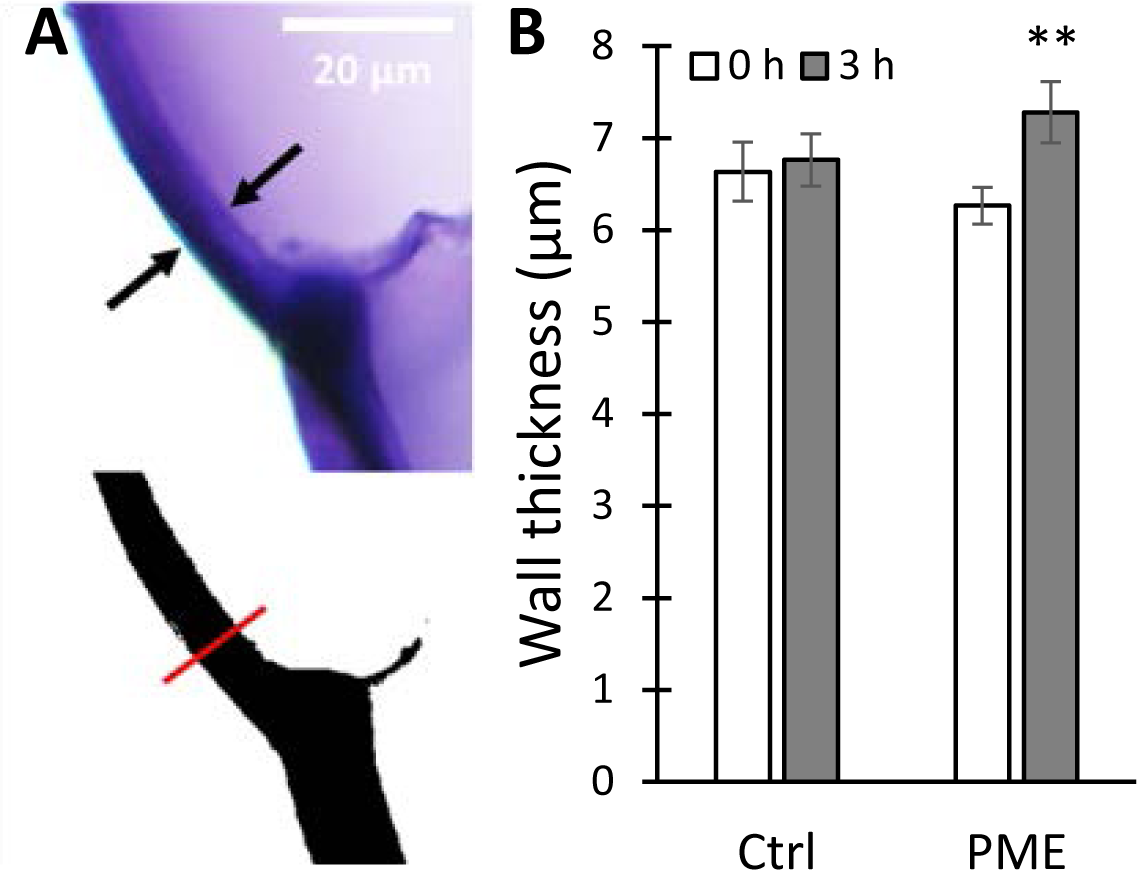
Cell wall swelling caused by PME treatment. (A) Onion scale cross-sections stained with toluidine blue; the wall thickness is marked by black arrows (top). Scale bar = 20 μm. Binary image was generated using ImageJ, red line shows the selection for wall thickness measurement (bottom). (B) Wall thickness comparisons at 0 h and 3 h, ± PME treatment. Values are means ± SEM (n = 20). Student’s t-test (paired, two-tail) was used to assess statistical significance (**-*p* < 0.01).

### PME treatment reduces cellulose microfibril resolution by AFM imaging

To assess the PME effect on the arrangement of cellulose microfibrils, we used the AFM to image the same wall surface before and after PME treatment. Distinct microfibril features can usually be resolved by AFM on the surface of native onion epidermal walls (Zhang *et al*., 2014; Zhang *et al*., 2016). After PME treatment, microfibril features were blurred or partially masked (Fig. 7), whereas treatment with BSA as a negative control did not obscure microfibrils (Supplemental Fig. S1). The blurring effect by PME may be the result of swollen HG chains near and between individual and bundled microfibrils, causing the AFM tip to glide over the surface without dropping into the spaces between microfibrils. Increased interaction of de-esterified HG with cellulose surfaces (Phyo *et al*., 2017) may also contribute to this effect. To confirm these visual impressions, we measured surface roughness of AFM height maps before and after PME treatment. Surface roughness, as measured by peak density, was reduced by 47% as a result of PME treatment (Supplementary Fig. S1) whereas BSA had no significant effect. These results confirm the visual impression of increased surface smoothness and poorer microfibril visibility after PME treatment.

**Fig. 7.**
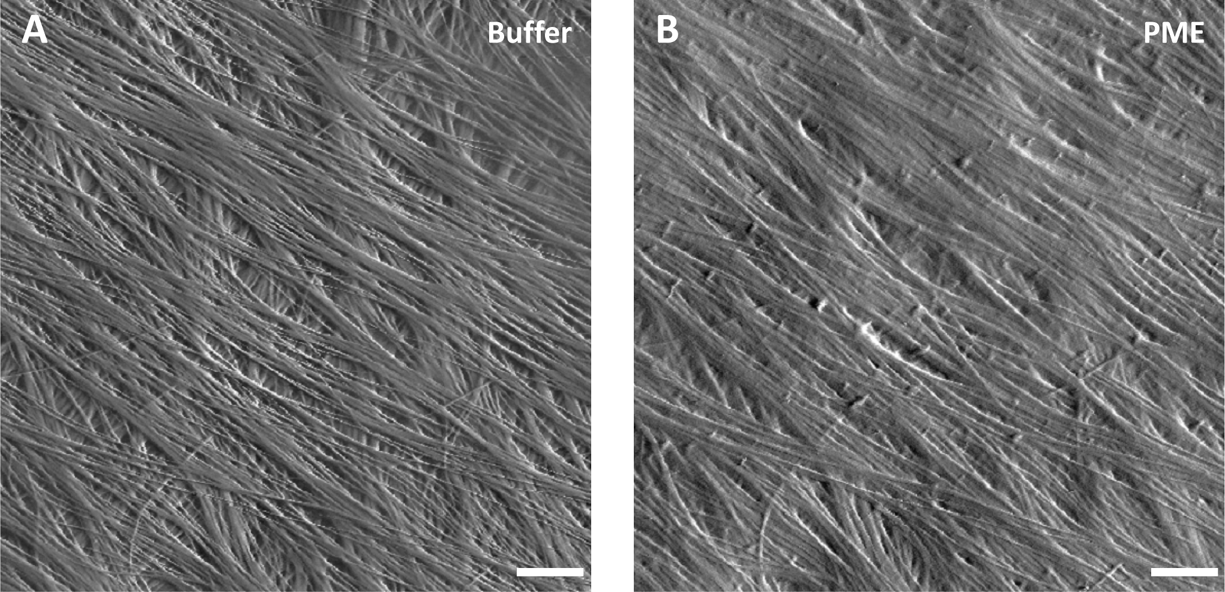
PME treatment reduces cellulose microfibril resolution by AFM imaging. (A) Peak force error image of onion epidermal wall surface in HEPES buffer. Scale bar = 20 nm. Distinct cellulose microfibrils and bundles well resolved by AFM. (B) Same area scanned after PME treatment. The resolution of cellulose microfibrils is reduced while fibrils in the underlying lamella were obscured. Similar results observed with six biological replicates.

### PME does not induce cell wall loosening

Creep and stress relaxation assays were used to assess PME’s ability to induce cell wall loosening. The stress relaxation assay measures the time-dependent decay in wall stress after the wall is extended and then held at constant length, whereas the creep assay measures the time-dependent increase in length when the wall is clamped at constant tensile force (Cosgrove, 2016). In stress relaxation assays of onion epidermal walls, acidic pH, which activates the α-expansins, enabled faster stress relaxation (Fig. 8A), whereas PME treatment resulted in a negligible change in stress relaxation (Fig. 8B). Thus PME treatment did not enhance wall stress relaxation.

**Fig. 8.**
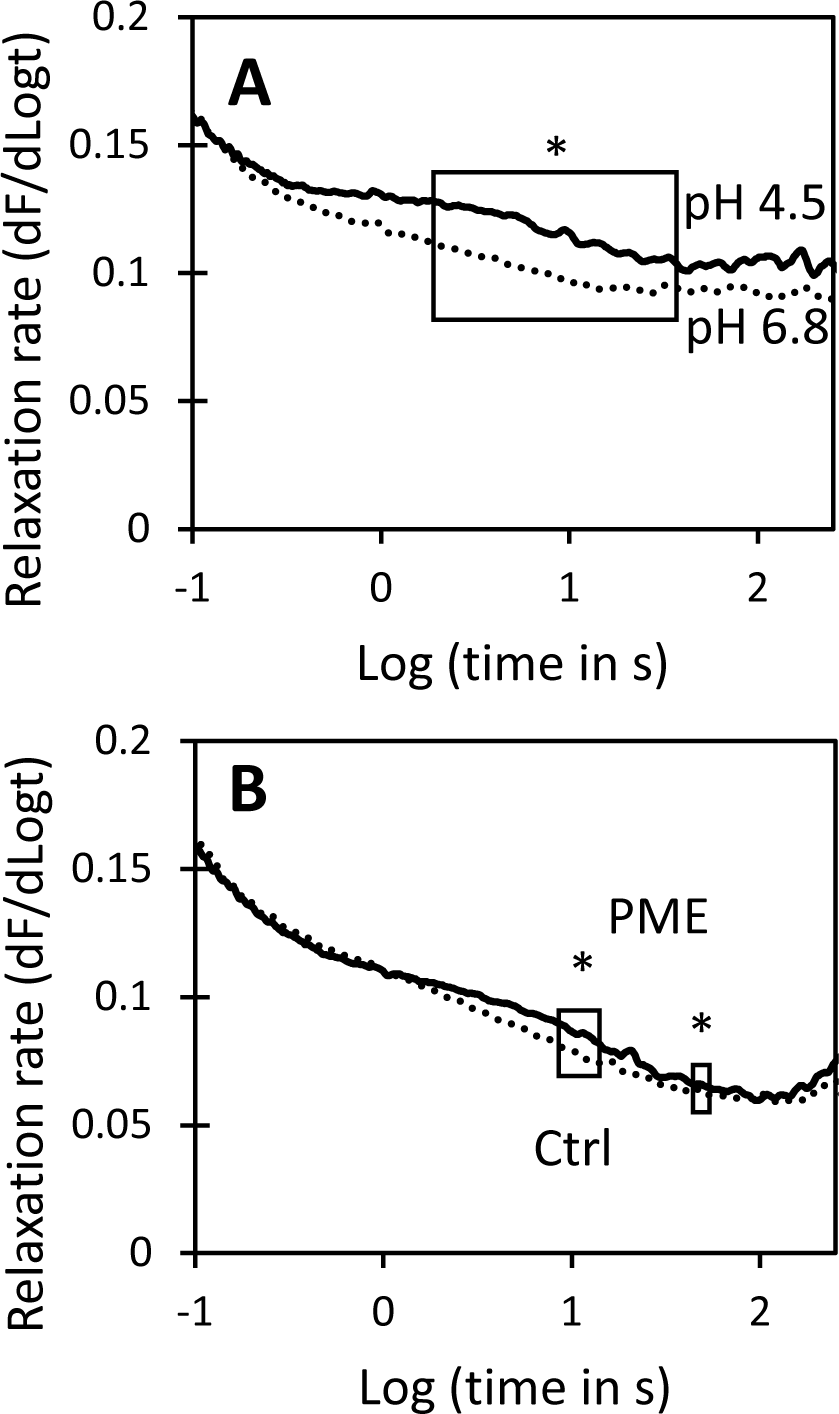
Effect of acid pH and PME pretreatment on stress relaxation spectra of onion epidermal walls. Each curve is the average of 7-8 data sets. Boxed regions show statistically significant difference in relaxation rate. Student’s t-test was used to evaluate statistical significance (* boxed region different at *p* < 0.05). (A) Walls in acidic (pH 4.5) buffer have greater stress relaxation than walls in neutral (pH 6.8) buffer. (B) Stress relaxation spectra of walls after 3 h incubation in 20 mM HEPES pH 7.5 ± PME.

For the creep experiments, wall strips were clamped at 0.1 N tension in neutral buffer and after the length stabilized the buffer was swapped for one containing PME. Length remained nearly constant for the duration of the experiment (90 min) and was not increased by PME addition (Fig. 9A). Thus we did not find evidence of PME-mediated wall loosening in this chemorheological creep assay.

**Fig. 9.**
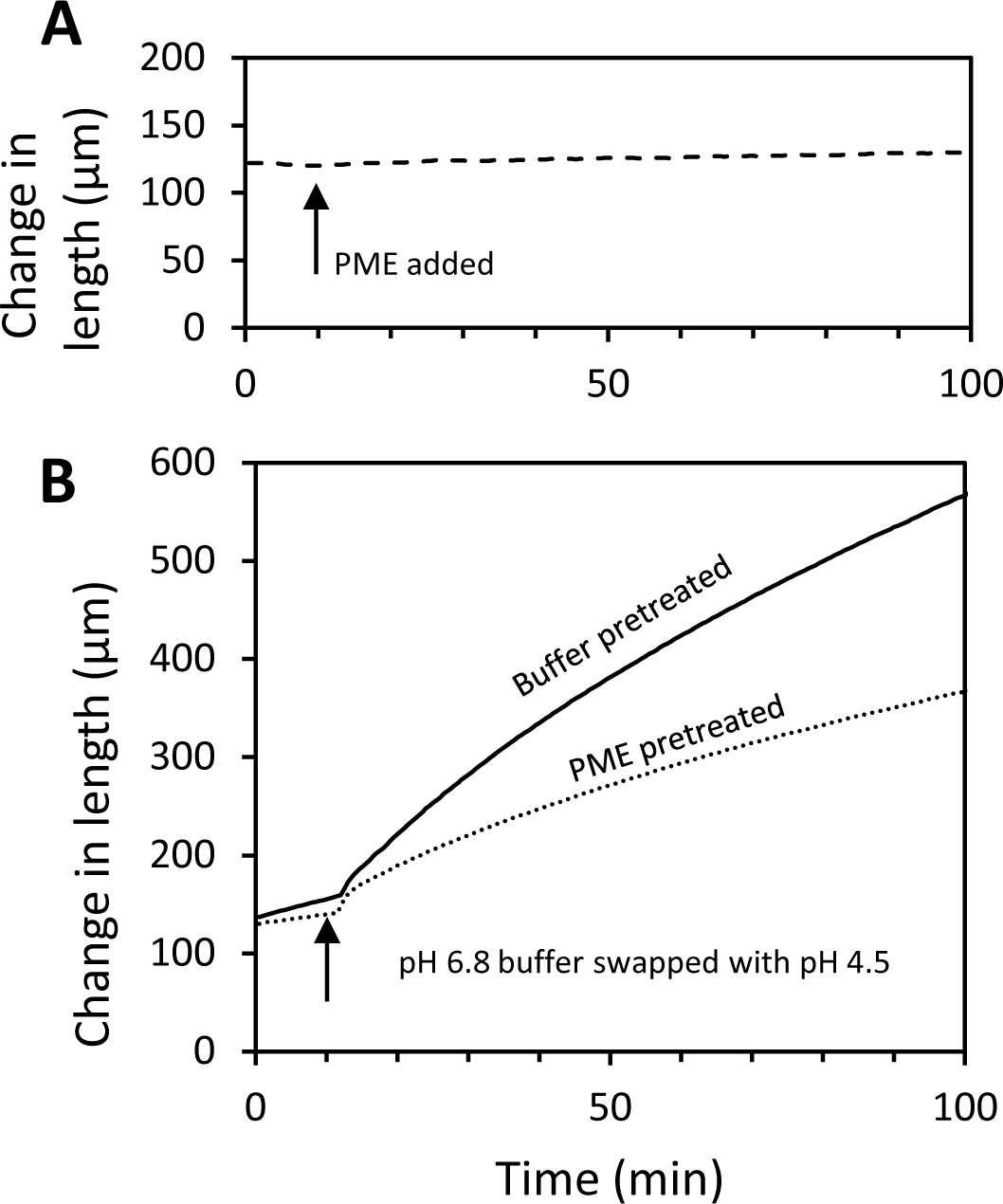
Influence of PME treatment on cell wall creep. (A) Addition of 100 μg/mL PME in pH 7.5 HEPES buffer does not induce creep. Walls were incubated in pH 7.5 buffer while clamped on the constant force extensometer. At the time indicated by the arrow, the buffer was exchanged with fresh buffer containing PME. (B) Acid-induced creep of walls pretreated for 16 h with buffer (solid) is substantially greater than creep of walls pretreated for 16 h with PME (dash). Curves are averages of 6 (PME) and 9 (buffer- and PME-pretreated) replicates.

We also tested whether PME pretreatment affected acid-induced extension, mediated by endogenous α-expansins. Wall strips, which were treated with hot methanol to inactivate endogenous PME and other enzymes but not endogenous α-expansins (McQueen-Mason *et al*., 1992), were pre-incubated in neutral buffer ± PME, then clamped at neutral pH in the constant-force extensometer, and tested for acid-induced extension by exchanging the neutral buffer for pH 4.5 buffer. The walls extended rapidly in acidic buffer. Sustained creep was diminished by ∼50% in the PME-treated walls compared with buffer-treated controls (Fig. 9B). Thus, despite the increase of wall hydration, tensile plasticity, and nanoindentation depth after PME treatment, expansin-mediated creep was reduced. We conclude that PME may selectively soften the epidermal wall under calcium-limited conditions, but we found no evidence for wall loosening by PME.

## Discussion

As detailed in the introduction, this study was initiated to resolve some of the confusion and speculation surrounding the action of PME on cell wall stiffness and extensibility (Levesque-Tremblay *et al*., 2015; Peaucelle *et al*., 2008). Our results show that – even in our simplified, cell-free system (isolated outer epidermal walls from onion) – a more nuanced appreciation of the complexity of wall biomechanics and the action of PME is needed to unpack this issue. Thus, measured by nanoindentation (Fig. 2) or by tensile plasticity (Fig. 3), PME softened the onion cell wall, yet it did not change tensile elasticity nor did it loosen the wall, assayed as the ability to induce cell wall creep (Fig. 9). Indeed, despite its selective softening and hydrating actions,

PME treatment actually hindered the wall loosening activity of endogenous α-expansins to induce acid-dependent extension. The wall became less extensible despite its increased hydration and plasticity. Thus, in this case plasticity is not a good predictor of the ability of the cell wall to undergo chemorheological creep – a conclusion similar to that of another recent study that employed other enzyme treatments (Zhang *et al*., 2019). These PME effects occurred without addition of calcium to the wall, so we conclude they were direct physico-chemical effects of HG de-esterification within the cell wall. Because endogenous wall enzymes were inactivated by pretreatment in hot methanol, the softening action did not involve HG degradation by endogenous pectinases or pectolyases. Hence, our answer to the question “How does PME affect wall biomechanical properties?” depends on the specific assay and requires the clarification that wall stiffness and wall creep are not tightly coupled (Cosgrove, 2018a; Zhang *et al*., 2019).

The results of the current study are relevant to understanding: (1) PME effects on wall mechanics, (2) the relationships of different biomechanical assays to each other and to growth, and (3) the relationships between wall structure and various biomechanical properties. These three points are discussed below.

### PME effects on wall mechanics

After PME treatment, the electrostatic potential of the onion wall (measured as zeta potential) became more negative, as expected for an enzyme that unmasks carboxylate groups of methylesterified HG (Moustacas *et al*., 1986). It is likely that cell wall swelling, hence greater wall hydration, after PME treatment resulted from increased electrostatic repulsion of negatively-charged HG chains (MacDougall *et al*., 2001; Ryden *et al*., 2000) and these effects in turn resulted in softening action measurable in the indentation and tensile tests. This later point is supported by the fact that MgCl_2_, which was used to reduce electrostatic fields within the wall, largely negated the PME effect on plasticity (Fig. 4) and indentation (Supplemental Table S4).

This scenario is consistent with previous work showing charge-dependent swelling of isolated pectins and tomato cell walls (MacDougall *et al*., 2001; Ryden *et al*., 2000; Zsivanovits *et al*., 2004), and supports the concept that pectin hydration influences wall thickness (Jarvis, 1992).

Hydration also influences wall extensibility in some conditions. For instance, wall dehydration by polyethylene glycol reduced wall extensibility in two studies (Edelmann, 1995; Evered *et al*., 2007). However, in the current study PME-mediated increase in wall hydration was associated with reduced cell wall creep, not higher creep, despite an increase in wall plasticity. Perhaps the increased electrostatic charge in the wall interferes with expansin-mediated creep (Wang *et al*., 2013), despite higher hydration. There are many potential mechanisms for such interference (Ricard, 1987). A more detailed look at the effects of electrostatic charge on cell wall rheology might be given insights into the basis of this charge effect.

The greater electrostatic charge after PME treatment amplified the sensitivity of the cell wall to exogenous calcium. For instance, calcium addition had little effect on tensile compliances (elastic or plastic) of buffer-treated walls, whereas after PME-treatment calcium addition substantially reduced wall plasticity (but not elasticity) (Fig. 4). These observations suggest the possibility that newly unmasked carboxylate groups participated in calcium crosslinking of HG, e.g. via the ‘egg box’ model (John *et al*., 2019; Morris *et al*., 1982), stiffening the matrix. However, the lack of effect on elasticity runs counter to this simple explanation, as more crosslinking of HG might be expected to reduce the elastic compliance, if the wall behaved like a fiber-reinforced hydrogel, as often assumed (Milani *et al*., 2013). Evidently a different model of the cell wall is needed to account for its complex and nonintuitive biomechanical behaviors (Zhang *et al*., 2019).

Other physical mechanisms may contribute to the reduced plasticity of PME-treated walls incubated with CaCl_2_: approximately half of the calcium effect may due to electrostatic shielding, judging from the effect of MgCl_2_ (Fig. 4) and assuming that Mg^2+^ does not form HG crosslinks (Thibault and Rinaudo, 1986); calcium-HG interactions may also reduce HG-cellulose interactions (Lopez-Sanchez *et al*., 2020; Phyo *et al*., 2017; Wang *et al*., 2015). Such interactions appear to be extensive, but their significance for wall mechanics is uncertain. A molecular understanding of the nature of cell wall plasticity, elasticity and the physical interactions between cell wall polymers is required to assess the relative contributions of these different biophysical mechanisms to wall mechanics.

Because the experiments in the current study imposed relatively large changes in HG methyl esterification (large in the biological context) and used high calcium concentrations, these results should be considered the extremes of possible PME-dependent and calcium-dependent changes in wall biomechanics. Whether similar changes occur *in vivo* is uncertain at this time.

Nevertheless, the nanoindentation results do offer a potential explanation for correlations between HG de-esterification and reduced indentation stiffness on Arabidopsis surfaces (Peaucelle *et al*., 2011; Peaucelle *et al*., 2015). How indentation stiffness relates to other wall properties is considered next.

### Relating different biomechanical assays to each other and to growth

One striking conclusion from this study is that different measures of wall biomechanics are not closely coupled to one another. Thus, PME action softened the wall, as measured by indentation and tensile plasticity, yet it did not result in wall loosening, as measured by cell wall creep. Wall creep is considered a fundamental mechanism of cell wall growth (Cosgrove, 2018a).

Consequently our *in-vitro* results thus do not support the concept that PME has direct wall loosening activity. Various indirect mechanisms for PME-mediated wall loosening have been proposed (Moustacas *et al*., 1986; Moustacas *et al*., 1991; Peaucelle *et al*., 2008) but they remain untested and wall loosening by PME remains unconfirmed.

These results with PME confirm and extend conclusions of another recent study likewise showing that wall softening and loosening are not tightly coupled in the onion epidermal wall (Zhang *et al*., 2019). In the case of onion epidermal walls, nanoindentation evidently does not serve as a reliable indication of tensile properties. Whether this is also true for other epidermal walls needs to be examined. Theory predicts that indentations with larger probes (1 μm or larger and at greater depths may be sensitive to wall tensile properties (Milani *et al*., 2013), but this prediction requires experimental validation.

### Relating wall structure to various biomechanical properties

Many conceptual depictions of the spatial arrangements and interactions of cellulose, hemicellulose and pectins in primary cell walls have been proposed since the 1970s, based largely on biochemistry and microscopy. These sketches make tacit inferences about wall mechanics, yet rarely has biomechanics been used to test the validity of these depictions, which can be viewed as graphical hypotheses in need of experimental testing. One such test by Park and Cosgrove (2012) rejected the concept that cellulose microfibrils were mechanically linked into a load-bearing network by xyloglucan tethers. Another concept advanced by Thompson (2005) imagines the wall to be a tangle of microfibrils whose ability to move are controlled by matrix viscosity and free volume between microfibrils. Results in the current study, as well as those in Zhang *et al*. (2019), seem at odds with this concept. Zhang *et al*. (2019) concluded that tensile (in-plane) properties (elastic compliance, plastic compliance, and creep) were largely determined by the network of laterally-connected cellulose microfibrils within individual lamellae of onion epidermal wall, whereas the indentation (out-of-plane) mechanics were largely controlled by pectins, along with some contributions from the cellulose microfibril networks (Zhang *et al*., 2019). Such results need to be incorporated into quantitative cell wall models that account for wall mechanics based on nanoscale structure and that provide both explanatory and predictive value (Smithers *et al*., 2019).

The current study of PME action indicates that electrostatics and hydration affect selective aspects of wall mechanics. Because PME does not cut the HG backbone, its biomechanical effects are likely the result of physical changes resulting from the increased negative charge on HG. This leads to charge repulsion of HG chains and swelling of the wall, which in turn affects the indentation properties. The increase in tensile plasticity may result from increased hydration, but higher electrostatic charge density within the wall may also influence polymer interactions directly. Despite PME-induced changes in nanoindentation, tensile elasticity was insensitive to HG esterification and to calcium cross linking. This is a remarkable result and suggests that static tensile forces are transmitted predominantly via the interconnected network of cellulose microfibrils with little mechanical contribution from HG networks. Other results by Zhang *et al*. (2019) point in the same direction.

### Concluding remarks

By use of isolated epidermal wall strips to explore the physical consequences of PME action, we avoided the complexity of living tissues, where cell anatomy and turgor pressure can complicate the interpretation of mechanical assays (Forouzesh *et al*., 2013; Weber *et al*., 2015) and where biological responses to pectin modifications can elicit far-ranging responses involving auxin, brassinosteroids, wall integrity sensors, and changes in transcription of thousands of genes (Braybrook and Peaucelle, 2013b; Wolf *et al*., 2012). Our results show that, in the absence of added calcium, PME softened the wall in the nanoindentation assay, potentially accounting for some previous AFM-based reports of wall softening associated with regions of HG de-esterification (Peaucelle *et al*., 2011; Peaucelle *et al*., 2015). A concomitant, though small, increase in onion wall plasticity, however, did not translate into a more extensible cell wall, as measured by cell wall creep, and so is unlikely to account for increased growth associated with regions of de-esterified HG. Wall biomechanics is multifaceted, nuanced, and offers a rich path for gaining insights into the hierarchical organization of cell wall polymers and the structural basis for wall plasticity, elasticity and other biomechanical properties.

## Supplementary data

Tables S1-4: Effects of 100 mM CaCl2 and MgCl2 on indentation of onion walls Fig. S1. Effects of PME and BSA on onion wall surface texture.

Data is available from the corresponding author upon request.

## Acknowledgments

This work was supported as part of the Center for Lignocellulose Structure and Formation, an Energy Frontier Research Center funded by the US Department of Energy, Office of Science, Basic Energy Sciences under award no. DE-SC0001090.

---

**Fig. S1.**
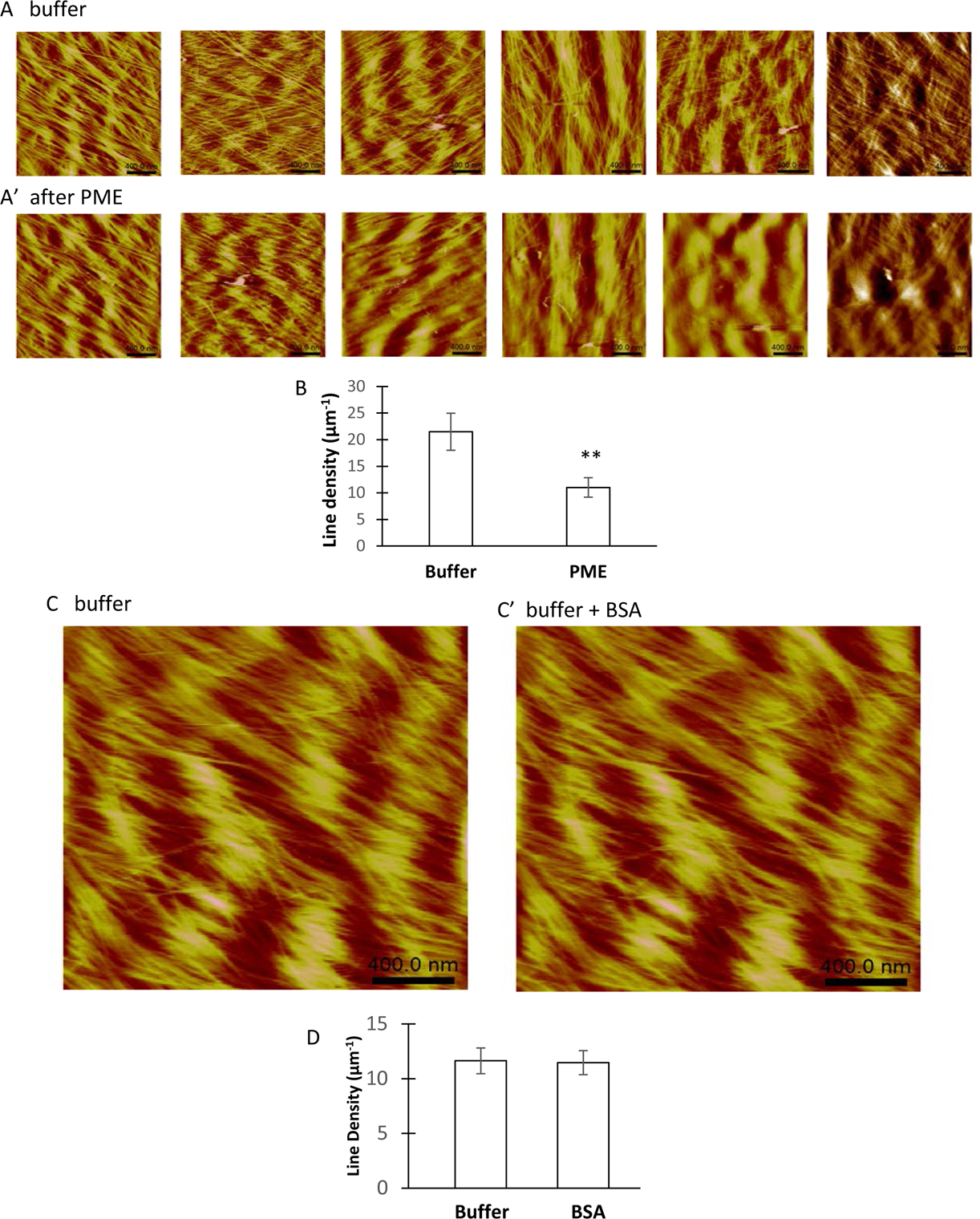
Effects of PME and BSA on onion wall surface roughness. (A) AFM height images of onion epidermal cell wall surface scanned in HEPES buffer. (A’) AFM images of the same cell wall sample scanned after PME treatment. Scale bar = 400 nm. (B) Roughness measurements of wall surface ± PME (n=6). Statistical significance was determined by Student’s t-test (paired, ** *p* < 0.01). (C) AFM height image of onion epidermal cell wall surface scanned in HEPES buffer. (C’) Corresponding AFM image of the same cell wall sample scanned with BSA in the buffer. Scale bar = 400 nm. (D) roughness measurements of wall surface ± BSA (n = 3). Statistical significance was determined by Student’s t-test (paired, ** *p* < 0.01).

**Supplementary Tables S1-4: Effects of 100 mM CaCl2 and MgCl2 on indentation modulus of onion epidermal walls with and without PME pretreatment. Values related to the cation effects are highlighted. Analysis of variance was used to factor out variablity due to the cell wall sample. For each treatment, ten indentations were measured in three cells.**

**Table S1.**
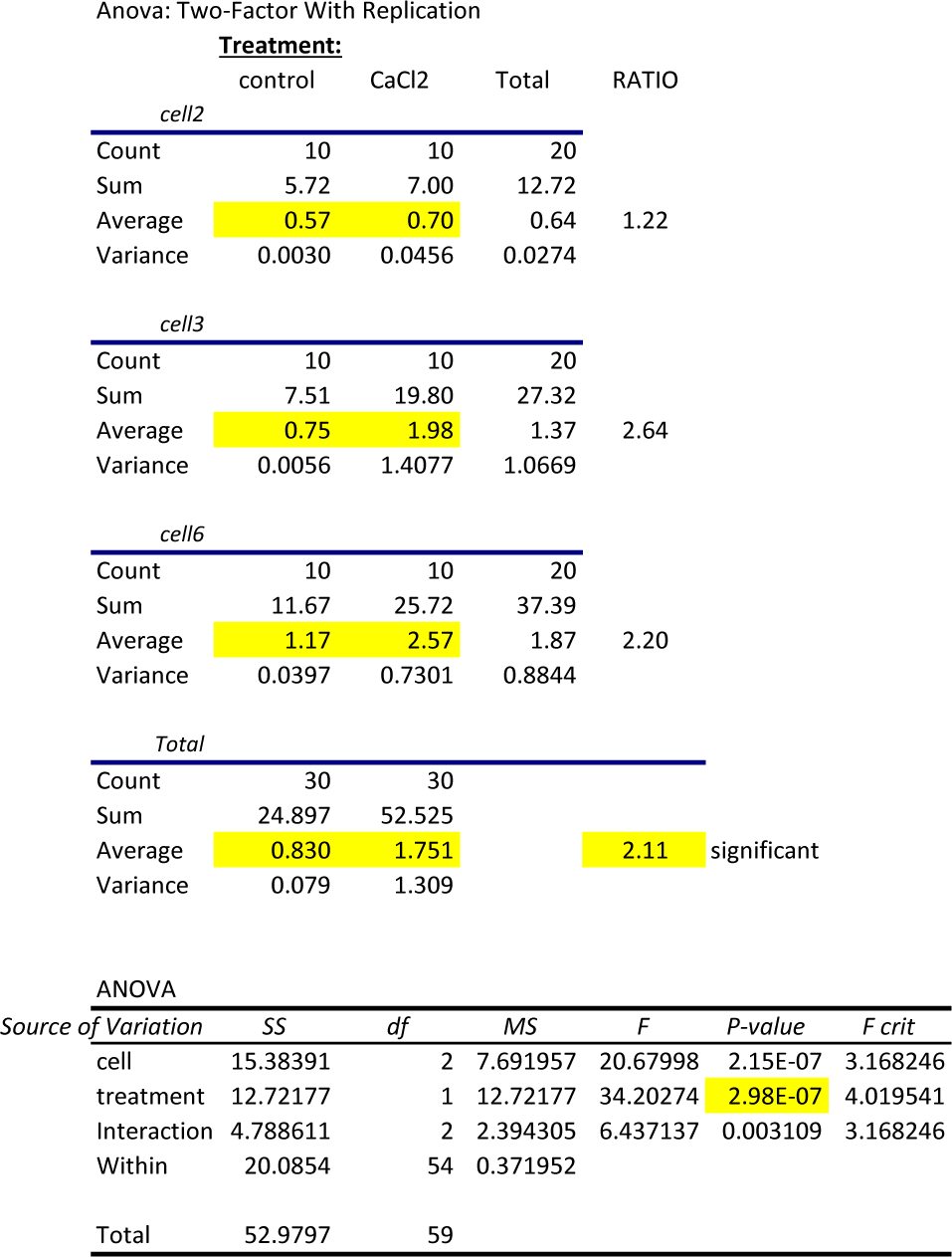
Effect of Ca on indentation modulus of untreated onion walls

**Table S2.**
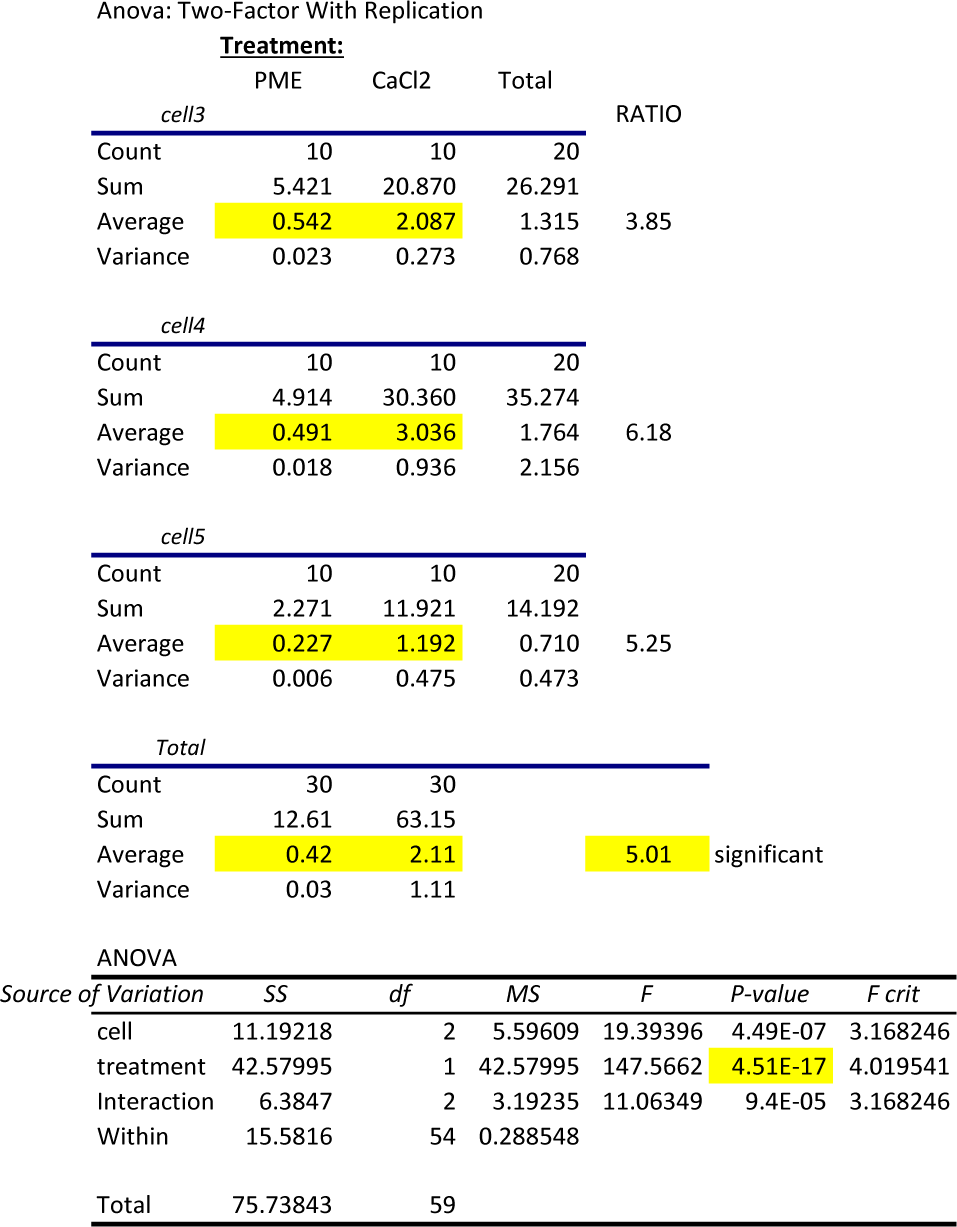
Effect of Ca on indentation modulus of PME-pretreated onion walls

**Table S3.**
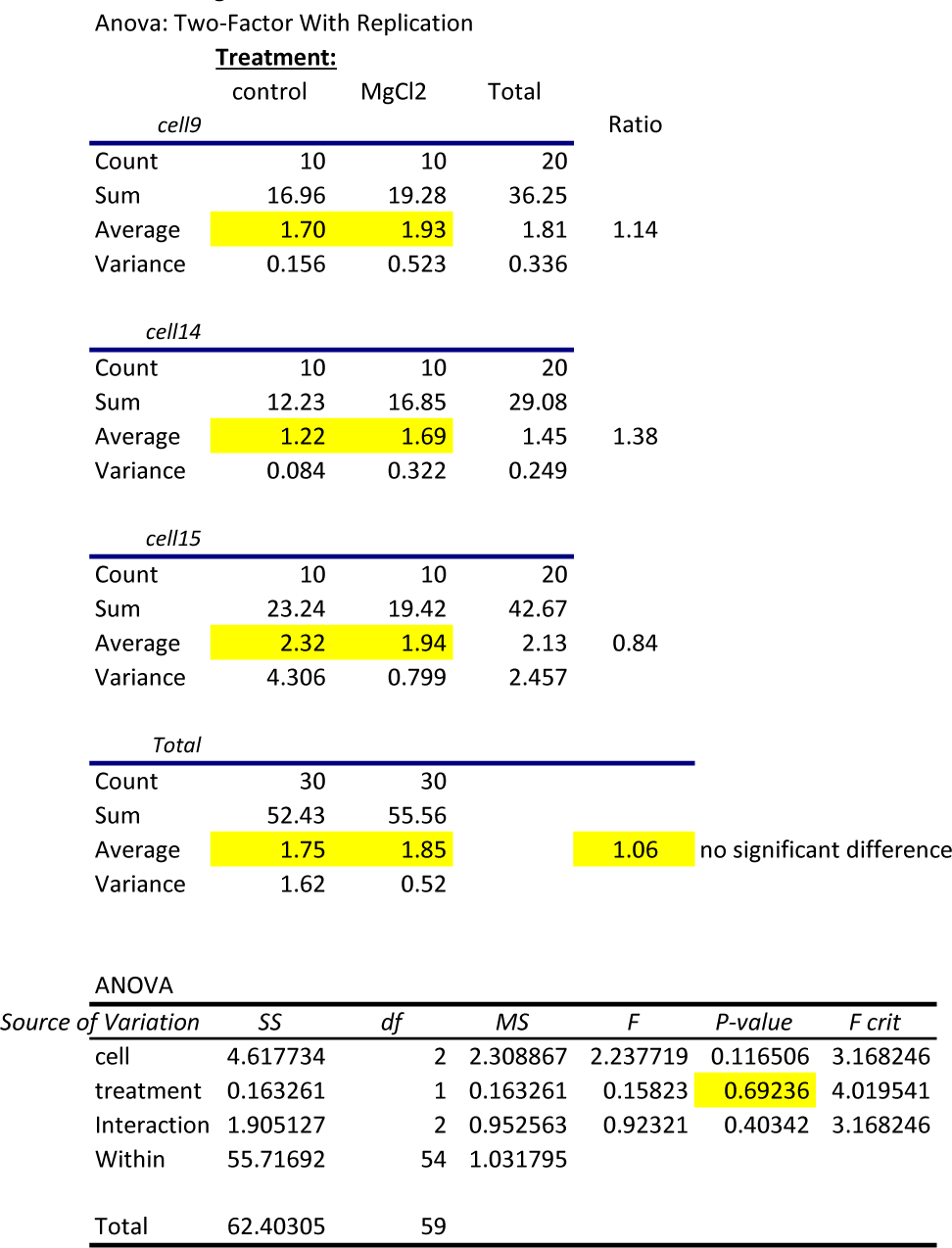
Effect of Mg on indentation modulus of untreated onion walls

**Table S4.**
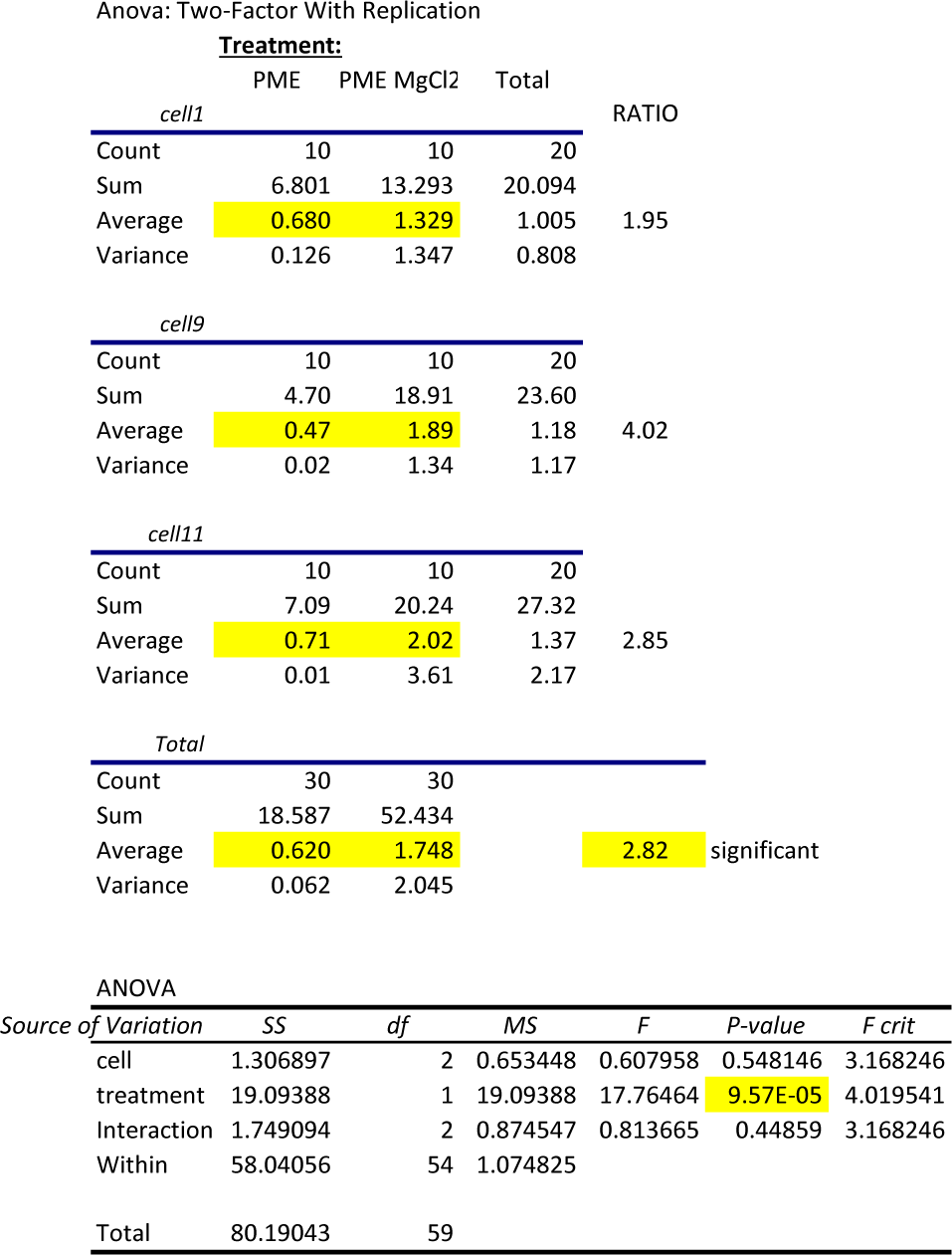
Effect of Mg on indentation modulus of PME-pretreated onion walls

